# Evaluation of Three Diets Based on Dry, Live, and Wet Feeds for Optimal Growth of Discus Fish (*Symphysodon* p.) Under Controlled Conditions in Florencia, Caquetá, Colombia

**DOI:** 10.1101/2024.12.23.630134

**Authors:** Alejandro Navarro-Morales, Jenniffer Tatiana Díaz Cháux, Helman Castañeda Castañeda, Diego Huseth Ruiz-Valderrama, Alexander Velasquez Valencia

## Abstract

This study evaluated the effect of three diets, wet (PAP), live (DAP) and commercial feed (COC), on the growth of juvenile discus fish (*Symphysodon* sp), under controlled environmental conditions in the Colombian Amazon. The PAP diet was made with animal protein and Amazonian fruits. The DAP diet consisted of the provision of live food (*Moina macrocopa*); Both diets were mixed in equal parts with the concentrate to be supplied to the fish. In the COC diet, only the commercial feed was supplied. The daily supply of each diet corresponded to 6% of the average body weight of the juveniles, adjusting the quantity in each measurement. Each treatment had four repetitions (aquarium), with a density of 0.32 individuals per liter for a total of 192 discus fish. The growth of the juveniles was monitored by taking morphometry every nine days for 63 days. The morphometric variables presented differences between the diets, the highest values were recorded in PAP and DAP. The morphometric variables such as total length and weight were positively associated with nutritional contents such as proteins, lipids and carbohydrates present in DAP and PAP. These components influenced the growth of the fish and were related to the feed conversion factor. It is suggested that fish fed with DAP and PAP continued to grow thanks to the biological and enzymatic stimuli provided by these foods, unlike the COC feeding with the lower growth reported. These results demonstrate that the combination of live foods prepared according to the habits and nutritional requirements of the species with concentrates promote the growth of the fish, promoting optimal development and general well-being of the discus fish during growth under controlled conditions.

## Introduction

The ornamental fish trade has increased significantly over the past decade (Dey, 2016), being recognized as a profitable economic activity and a driver of employment worldwide (Evers et al., 2019). Colombia ranks among the top ten countries in the global trade of wild ornamental fish (Dey, 2016). This is due to its great environmental, geological, and hydrological diversity (Rangel-Ch, 2015), which supports a high number of species with ornamental potential in this country (Sánchez-Páez & Muños-Torres, 2015). Among these are fish of the order Cichliformes, characterized by their variety of colors, body shapes, and behaviors. Species such as the Banded cichlid (*Heros severum*), Demon eartheater (*Satanoperca jurupari*), Angelfish (*Pterophyllum scalare*), and Discus (*Symphysodon aequifasciatus*) (Ureña et al., 2014; Aguinaga et al., 2015; Dey, 2016; Evers et al., 2019; Ortega et al., 2021) are among the most traded at national and international scales (Kullander & Silfvergrip, 1991; Ortega et al., 2021).

Most ornamental fish for commercial purposes are extracted from their natural environments, which has led to a decline in wild populations of some species (Moncelano & Franco-Ortega, 2020). However, certain species, such as the Discus fish (*Symphysodon aequifasciatus*), are being bred for commercial purposes under controlled conditions by breeders using empirical and artisanal methods (Jiménez-Rojas et al., 2012; Pava-Escobar et al., 2022; Navarro-Morales et al., 2023). The lack of technical knowledge, such as the formulation of specific diets tailored to the species’ biology, has resulted in high mortality rates and low productivity (Silva, 2018; López-Rodríguez et al., 2019; Hodar et al., 2020).

The Discus fish is characterized by a short and robust lower jaw, bicuspid teeth, a small stomach, and long intestines (Crampton, 2008; Burress, 2015). These features indicate an omnivorous diet composed mainly of plant material and small invertebrates (Zavala-Camin, 1996), in contrast to other cichlid species with piscivorous or insectivorous diets associated with more developed (i.e., larger) stomachs (Burress et al., 2015).

Many breeders and hobbyists are unaware of the feeding habits (omnivorous), digestive morphology, and physiology of this species (Blom & Dabrowski, 2000; Chong et al., 2003; Ortiz, 2010). Consequently, they often provide any type of feed (e.g., commercial fish feed designed for consumption fish) that lacks appropriate nutritional value (Sales & Janssens, 2013; Stevens et al., 2017) and the necessary supplements for optimal development (calories, lipids, proteins, among others) (NRC, 2011; Wen et al., 2018; Velasco-Garzón & Gutiérrez-Espinosa, 2019). Inadequate feeding reduces growth, increases the proliferation of diseases, and raises mortality rates in both juveniles and adult breeders (Jiménez-Rojas et al., 2012; Millan & Perez, 2017). As a result, optimal productivity is compromised, leading to significant economic losses for breeders (Velasco-Santamaria & Corredor-Santamaria, 2011).

Despite these nutritional challenges, few studies have explored the optimal diets for Discus fish under controlled production conditions (Navarro-Morales et al., 2023). Alternatives such as live feed cultivation and the preparation of wet feeds, based on the biology of this species, have been proposed. Luna-Figuera et al. (2017) suggest that providing live feed such as Artemia (*Artemia franciscana*), water fleas (*Daphnia magna* – *Moina macrocopa*), and mosquito larvae (*Tubifex tubifex*) contributes essential nutrients and stimulates predatory behavior (Velasco-Garzón & Gutiérrez-Espinosa, 2019). Wet feeds, prepared using various ingredients and formulations, have been shown to approximate the species’ nutritional requirements (Fuentes-Quesada et al., 2018).

In this context, the present study evaluated the growth response of juvenile Discus fish to three diets, comprising combinations of commercial feed, live feed, and wet feed prepared for *Symphysodon* sp. Additionally, the study assessed the organoleptic properties of the combined diets to determine which combinations contribute to optimal growth under controlled conditions in the Colombian Amazon region.

## Methodology

### Experimental Setup

The experimental phase of this study was conducted over three months, from August 24 to November 25, 2023, at the Biodiversity and Peace Research Center for the Amazon Region (CIBIPAZ) in Florencia, Caquetá, Colombia. A fully enclosed facility measuring 5 m in length, 3 m in width, and 2 m in height was constructed to control environmental variables (temperature, noise, and wind) that could affect the experiment’s outcomes. Inside the facility, two Samurai® 1500W heaters maintained a constant temperature of 28°C, while artificial lighting was provided for 12 hours daily. An air turbine, connected via a hose network to sponge filters, facilitated filtration and oxygenation in each aquarium.

Twelve 50 L substrate-free aquariums were used, maintaining a stocking density of 0.3 L/individual, equivalent to 16 45-day-old juveniles per aquarium, for a total of 192 Discus fish. These fish were varieties bred and commercially available in Colombia. According to Mattos et al. (2016), this volume is suitable for rearing and growing Discus fish (*Symphysodon* sp.). The average initial measurements of the juveniles were weight (1.46 ± 0.15 g), body length (33.8 ± 0.75 mm), and body width (17.3 ± 1.05 mm).

### Physicochemical Parameters

Maintaining optimal physicochemical parameters in the water minimizes stress and prevents disease outbreaks during the rearing of Discus fish (Pierhoen et al., 2012; Wen et al., 2017; Lewisch et al., 2018; Swain et al., 2020). Therefore, daily water changes of 80% of the total aquarium volume were performed (Ng et al., 2023), using chemically free water from a natural source. Monitoring of physical parameters, such as temperature (°C), total dissolved solids (TDS), and electrical conductivity (EC), was carried out using a BLE-C600® multiparameter digital device. For chemical parameters such as ammonium (NH4), nitrites (NO2), and pH, an API® titration kit was used. The physicochemical parameters in the aquariums remained constant and within ideal ranges for the fish: TDS (43.31 ± 1.22 ppm), EC (83.61 ± 1.17 μS/cm), NH4 (0.31 ± 0.09 ppm), NO2 (0.22 ± 0.02 ppm), pH (6.91 ± 0.50), and temperature (28.82 ± 0.96°C).

### Dietary Treatments

To evaluate the growth response to different diets in juvenile Discus fish, a completely randomized design was used with three treatments (diets) and four replicates each. Treatment 1 (DAP): Equal proportions of live feed (*Moina macrocopa*) and commercial feed. Treatment 2 (PAP): 50% wet feed and 50% commercial feed. Treatment 3 (COC): 100% commercial feed.

A daily ration equivalent to 6% of the juveniles’ body weight was provided, adjusted every nine days based on individual growth assessments. This ration was distributed across four feeding sessions, starting at 9:00 AM and ending at 6:00 PM, with three-hour intervals between feedings.

For live feed (*Moina macrocopa*), six containers were prepared with a mixture of 10 L of green algae (*Chlorella* sp.) and 100 g of water fleas, exposed to direct sunlight. A fermented nutrient compound was used to sustain the algae culture. The wet feed, or paste, was prepared from various ingredients based on the nutritional requirements of Discus fish (Santos et al., 2022). This included animal protein (beef, seafood, and fish), vegetables, and Amazonian fruits such as açaí (*Euterpe oleracea*), camu camu (*Myrciaria dubia*), and milpes (*Oenocarpus bataua*) in a 2:1 ratio. The mixture was frozen in hermetically sealed bags. For commercial feed, Discus Granules by SERA® were used. This soft feed contains a high proportion of vegetable protein, suitable for the digestive physiology of Discus fish. A proximal analysis and nutritional composition assessment were conducted in triplicate using bromatological tests for moisture, protein, fat, ash, fiber, and carbohydrates in the dietary treatments (Table 1).

**Table 1.** Nutritional composition of dry weight for diets provided to juvenile Discus Fish (*Symphysodon* sp).

| Nutritional Components | COC | DAP | PAP |
| --- | --- | --- | --- |
| Calories (kcal) | 383 | 61 | 237 |
| Total protein (g) | 54.10 | 4.50 | 8.10 |
| Total lipids (g) | 9.20 | 3.80 | 18.3 |
| Crude fiber (g) | 3.70 | 0.40 | 1.50 |
| Ash (g) | 7.37 | 3.94 | 7.89 |
| Carbohydrates (g) | 21.0 | 2.10 | 9.90 |
| Moisture | 5.0 | 85.26 | 54.31 |

COC, dry feed; DAP, live feed; PAP, wet feed; kcal, kilocalories; g, grams.

### Measurements

he growth of the Discus fish was monitored on eight occasions at nine-day intervals. Ten morphometric parameters were evaluated for each individual, including weight, total length (TL), body length (BL), head length (HL), trunk length (TL), body width (BW), mouth-to-eye length (ME), eye diameter (ED), dorsal fin length (DFL), body mass index (BMI), and shape index (SI). To take the measurements, the fish were individually removed from the aquariums. Each fish was placed in a Petri dish lined with graph paper (1 mm x 1 mm) on a precision analytical balance (0.001 g) to record weight and photographed using a Nikon® P1000 camera. The photographs were processed in CorelDrawX3 software to extract morphometric measurements. The defined measurement scale provided by the graph paper allowed for precise data collection while minimizing stress caused by handling, thereby preserving the health of the fish.

### Statistical Analysis

To determine significant differences in morphometric parameters among the treatments, an analysis of variance (ANOVA) was performed, with tests of normality and homoscedasticity conducted for each response variable. Any significant differences identified between treatments were analyzed further using Fisher’s LSD multiple comparison test (P < 0.05). These analyses were conducted using InfoStat software version 2020 (Di Rienzo et al., 2020).

To estimate weight gain relative to feed intake for each diet, the feed conversion ratio (FCR = feed intake * weight gain^−1^) was calculated. Feed consumption per individual Discus fish (*Symphysodon* sp.) was determined using the formula: (CA= feed intake*n^−1^*t^−1^) where n is the number of individuals and t is the duration in days. The relationship between feed consumption (CA) and total length (mm) was modeled using a nonlinear regression based on a power model; *CA*=*a*\* *L^b^*.

The proximal composition of the diets was included as dependent variables, while weight and total length were used as response variables. Treatment was treated as a random effect to account for variability, and z-transformed (centered and standardized) fixed effects were used to compare regression coefficients across models (Schielzeth, 2010).

The effects of proximal composition on weight and total length were analyzed using Bayesian linear mixed models (BLMM) with a Gaussian error distribution. The model included all fixed effects, and the variance inflation factor (VIF) was calculated for each variable. Variables with VIF > 2 were excluded to avoid multicollinearity, and the models were re-executed (Zuur et al., 2010). Consequently, total fat, ash, and carbohydrates were excluded from the final model. All analyses were performed in R version R 4.4.1 (R Development Core Team, 2021) using the ‘usdm’ package (Naimi, 2015) and Bayesian methods with default priors for fixed and random effects in the ‘MCMCglmm’ package (Hadfield, 2010). A minimum of 1,000 posterior distributions was obtained by running 13,000 iterations for each model, with a burn-in period of 1,000 iterations and a thinning interval of 3,000 iterations. Statistical significance was determined if the confidence intervals did not overlap zero. A Markov chain summary for each predictor’s intersection in the Bayesian linear mixed model was visualized.

## Results

Juvenile Discus fish (*Symphysodon* sp.) consumed the entirety of the daily rations provided in all three treatments throughout the experimental period. No differences were observed in the initial weight and body length measurements of juveniles among the PAP, DAP, and COC treatments (F = 0.60; df = 2; P = 0.5711). During the first three measurement intervals, individuals in all three diet groups exhibited rapid increases in size and weight. However, growth slowed during the fifth and sixth measurements, with no significant differences detected among treatments during this period (F = 0.12; df = 2; P = 0.8852). After the sixth measurement, individuals resumed growth in both length and weight, continuing until the end of the experiment (Fig 1).

**Figure 1.**
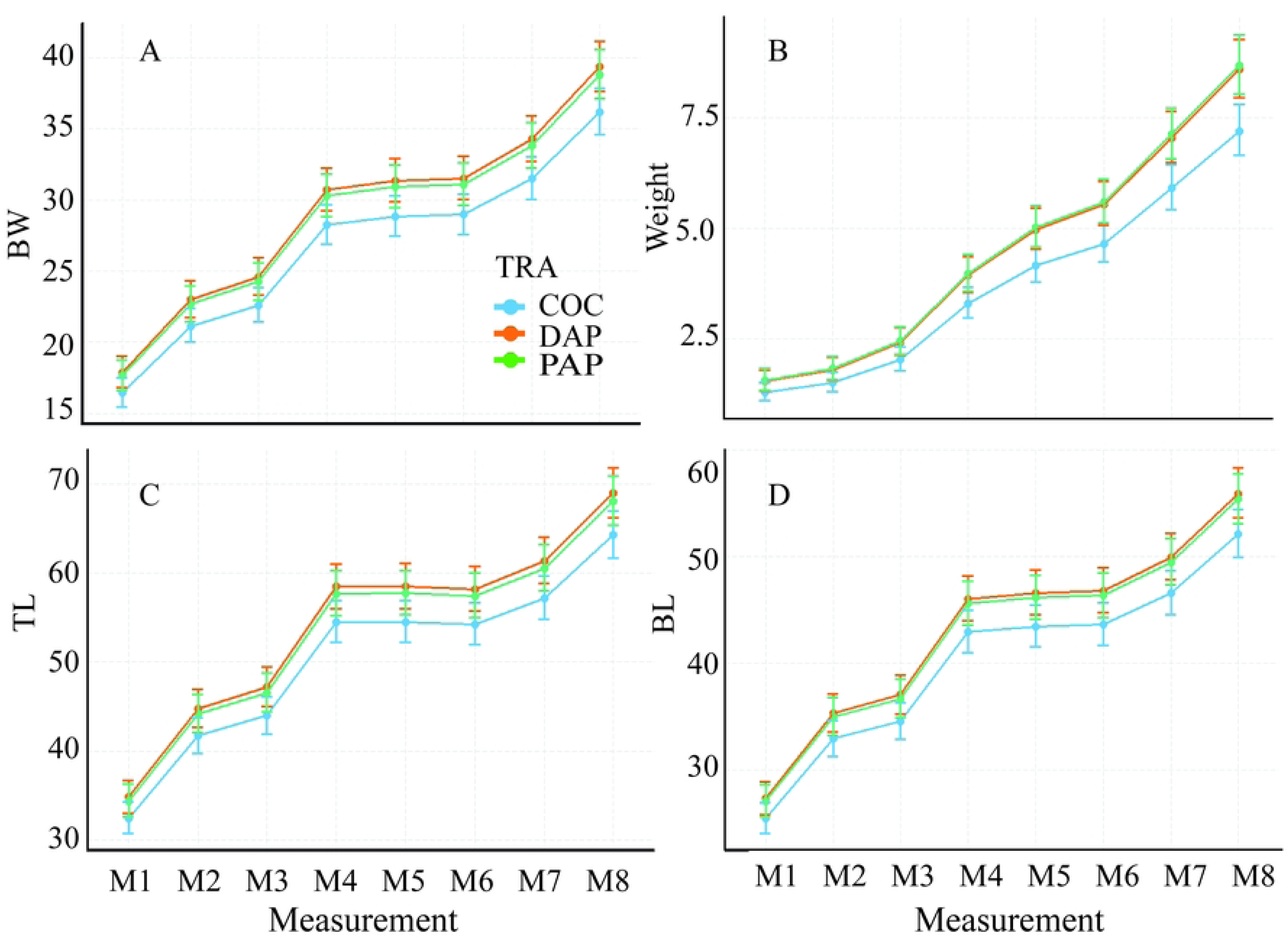
Morphometric growth curves for juvenile Discus fish (*Symphysodon* sp.) for the three diets. TRA: dietary treatments. COC: dry feed. DAP: live feed. PAP: wet feed. (A) BW: body width (mm). (B) Weight (g). (C) TL: total length (mm). (D) BL: body length (mm).

Growth in six length measurements and weight during the experimental period was greater in individuals fed with DAP and PAP diets (Fig 2). No significant differences were observed in eye diameter (ED), body width (BW), or mouth-to-eye length (ME) across treatments (P > 0.05), highlighting the effects of the diets on growth.

**Figure 2.**
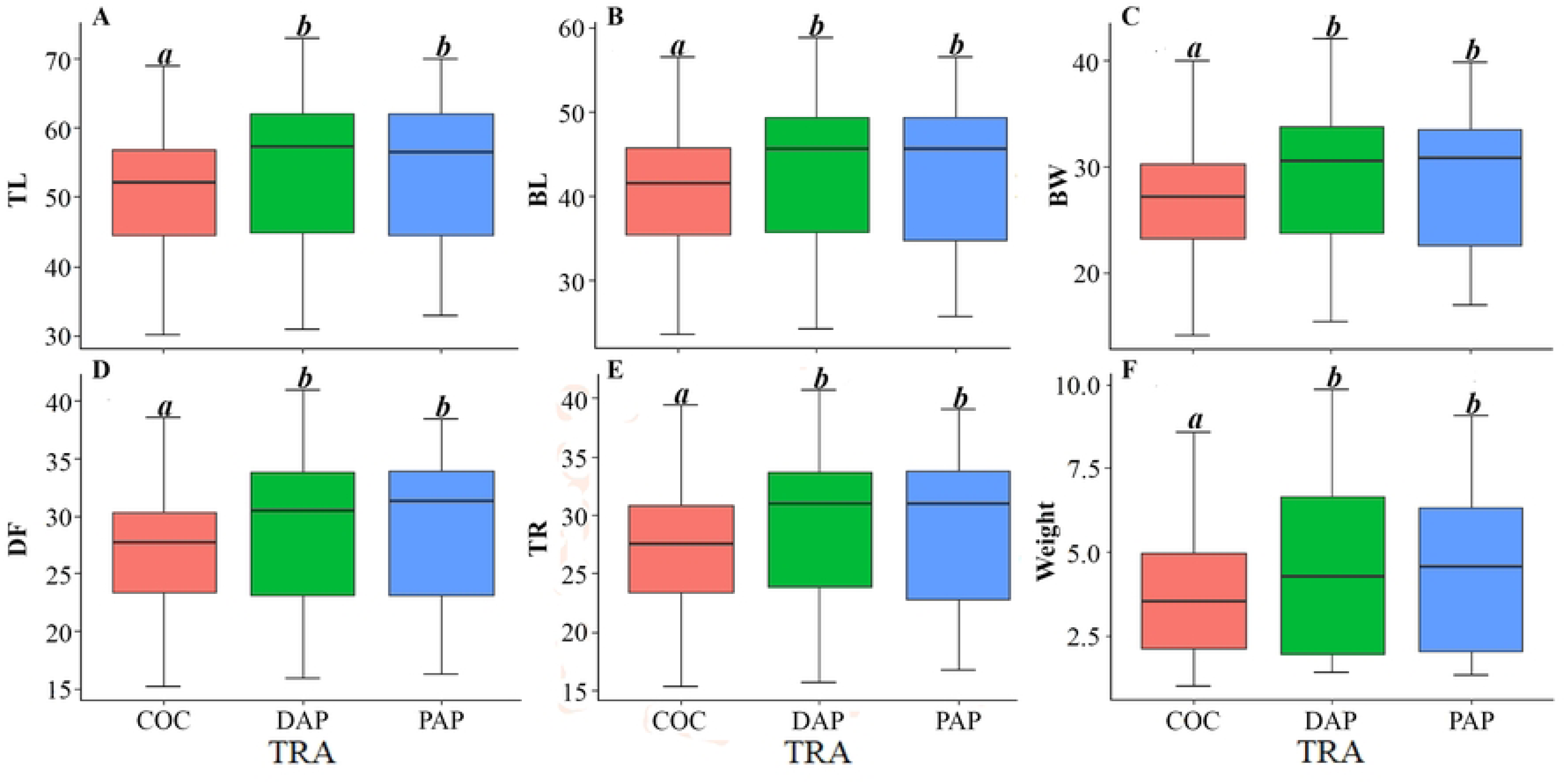
Boxplots of morphometric variables in juvenile Discus fish (*Symphysodon* sp.) for the three diets. TRA: dietary treatments. COC: dry feed. DAP: live feed. PAP: wet feed. (A) TL: total length (mm). (B) BL: body length (mm). (C) BW: body width (mm). (D) DF: dorsal fin length (mm). (E) TL: trunk length (mm). (F) Weight (g). Groups sharing the same letter are not significantly different (P > 0.05).

The feed conversion ratio (FCR) was highest in fish fed the COC diet (F = 3.40; df = 20; P = 0.0459). FCR values were initially high across all three diets but decreased toward the end of the experiment (Fig 3).

**Figure 3.**
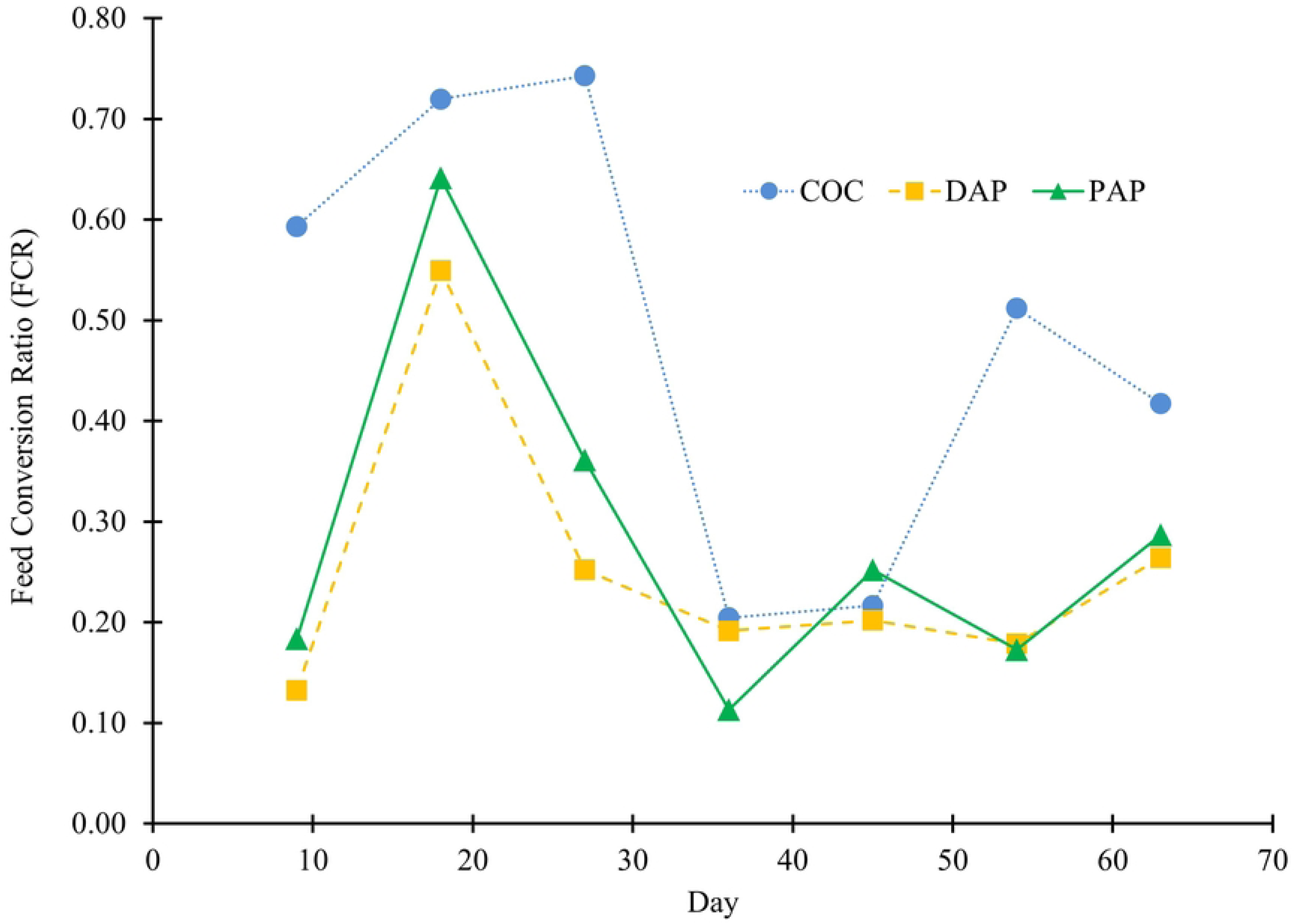
Feed conversion ratio for the three evaluated diets for juvenile Discus fish (*Symphysodon* sp.). COC: dry feed. DAP: live feed. PAP: wet feed. FCR: g feed*g fish^−1^.

The relationship between feed consumption (CA) and total length (TL) differed significantly across treatments (KW = 8.97; n = 21; P = 0.0112). Consequently, the potential curves for feed consumption were represented by three equations with different exponential values (Fig 4).

**Figure 4.**
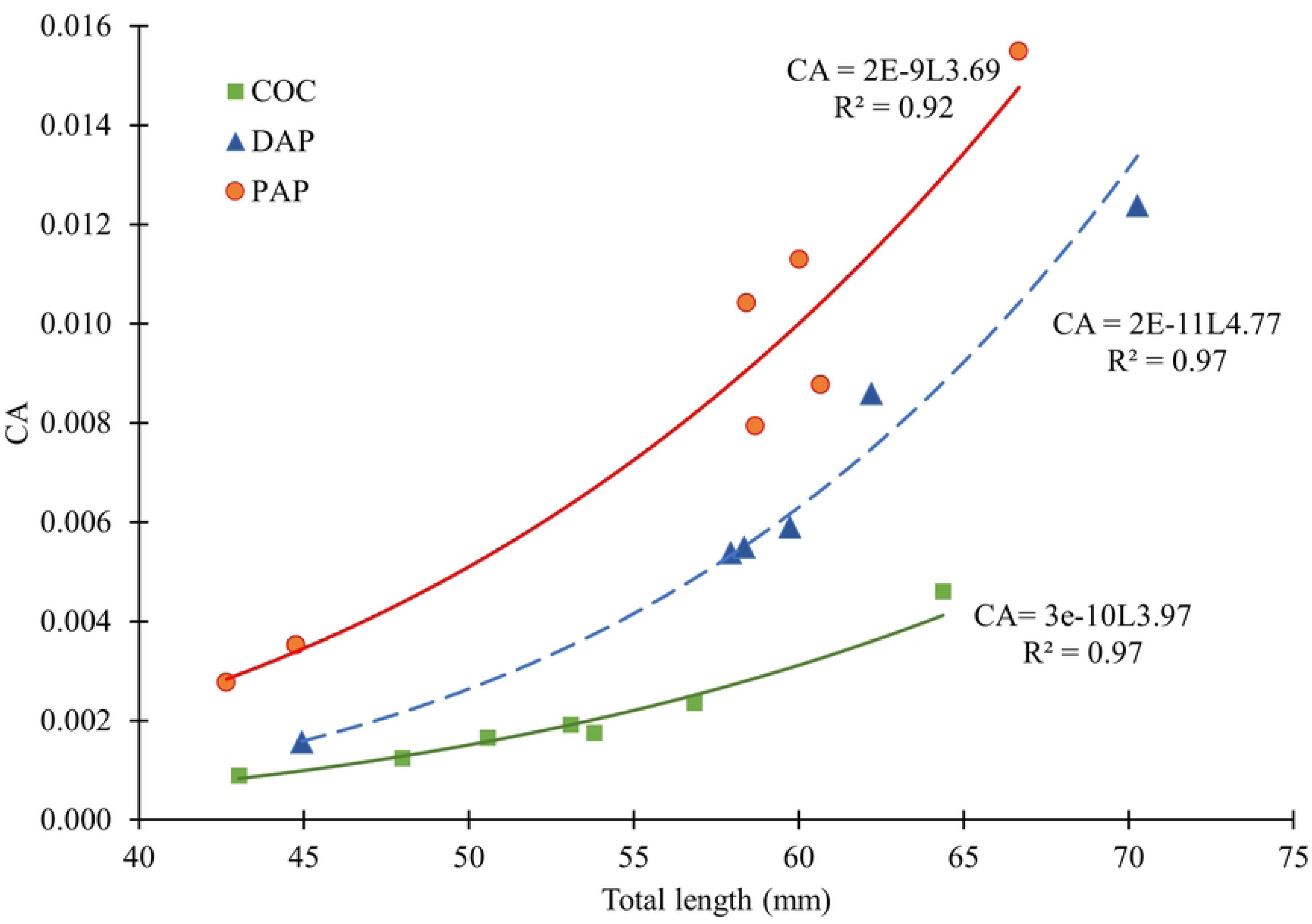
Relationship between the feed consumption and total length in juvenile Discus Fish (*Symphysodon* sp.). COC: dry feed. DAP: live feed. PAP: wet feed. CA: g feed*individual^−1^*day^−1^).

### Effects of Proximal and Bromatological Composition on Weight and Total Length

Weight and total length were positively influenced by protein concentration (pMCMC = 0.0029) and total fat content (pMCMC = 0.0033; pMCMC = 0.00667), exhibiting similar effects (Fig 5, Table 2). Conversely, carbohydrates, ash, and fiber did not significantly affect these morphometric parameters.

**Figure 5.**
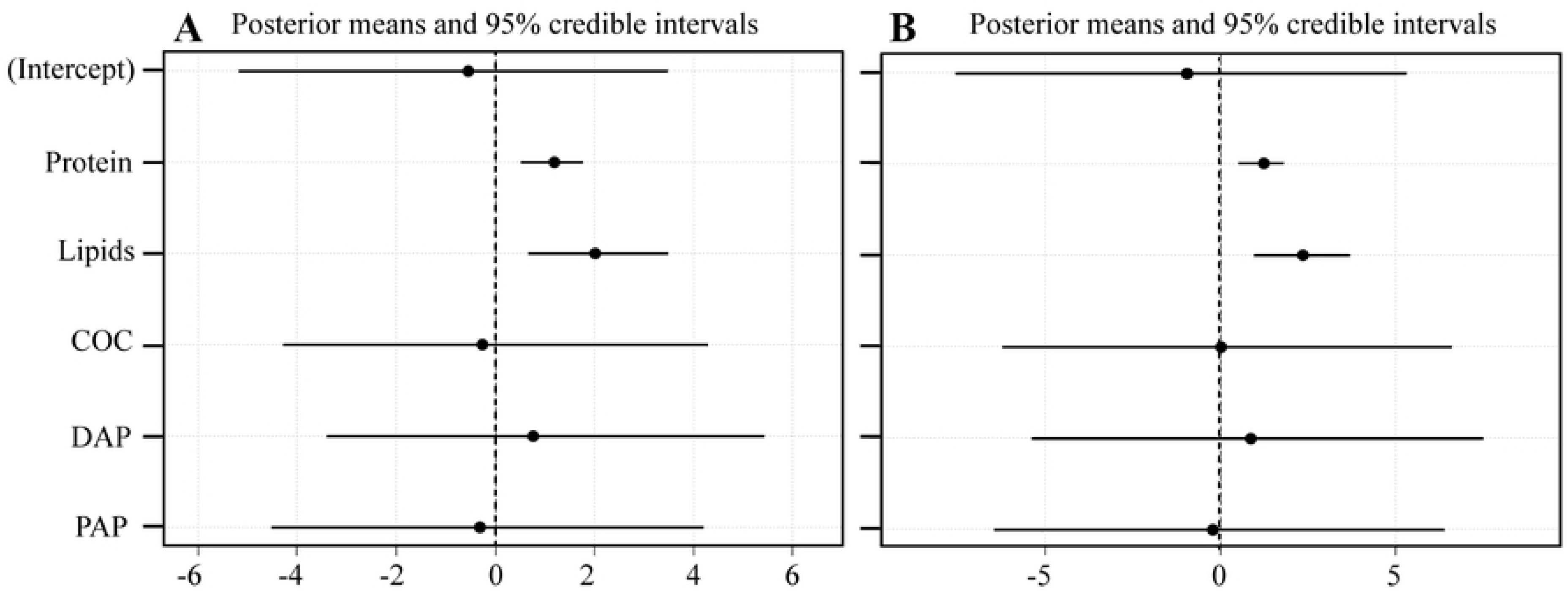
Standardized regression coefficients of BLMM in juvenile Discus fish (*Symphysodon* sp.). (A) total length (mm). (B) weight (g). BLMM: Big Linear Mixed Models. COC: dry feed. DAP: live feed. PAP: wet feed. TRA: dietary treatments. Total points indicate the posterior mean, and lines represent 95% confidence intervals. A variable is considered statistically significant if its confidence interval does not overlap zero.

**Table 2.** BLMM results for measurements, proximal and bromatological composition of diets supplied to juvenile Discus fish (*Symphysodon* sp.).

| Measurements | Components | Post mean | 95% CI | pMCMC |
| --- | --- | --- | --- | --- |
| Weight | Protein | 1.207 | 0.371 a 1.641 | 0.002 |
|  | Lipid | 2.235 | 0.842 a 3.171 | 0.011 |
| Total length | Protein | 1.082 | 0.547 a 1.641 | 0.002 |
|  | Lipid | 1.805 | 0.651 a 3.171 | 0.011 |

BLMM: Big Linear Mixed Models. CI: confidence intervals. pMCMC: Particle Markov chain Monte Carlo methods.

In the principal component analysis, the first principal component (PC1) explained 82.7% of the variability, while the second principal component (PC2) accounted for 10.2%. In PC1, morphometric variables were positively associated with the PAP diet and the contents of protein, fats, and carbohydrates. Accordingly, body lengths and weights of juvenile Discus fish were correlated with the high concentrations of proteins and fats in the PAP and DAP diets. However, PC2 grouped the composition of proteins, carbohydrates, and ash with the COC diet, and morphometric measurements were negatively associated with the DAP diet in this component (Fig 6).

**Figure 6.**
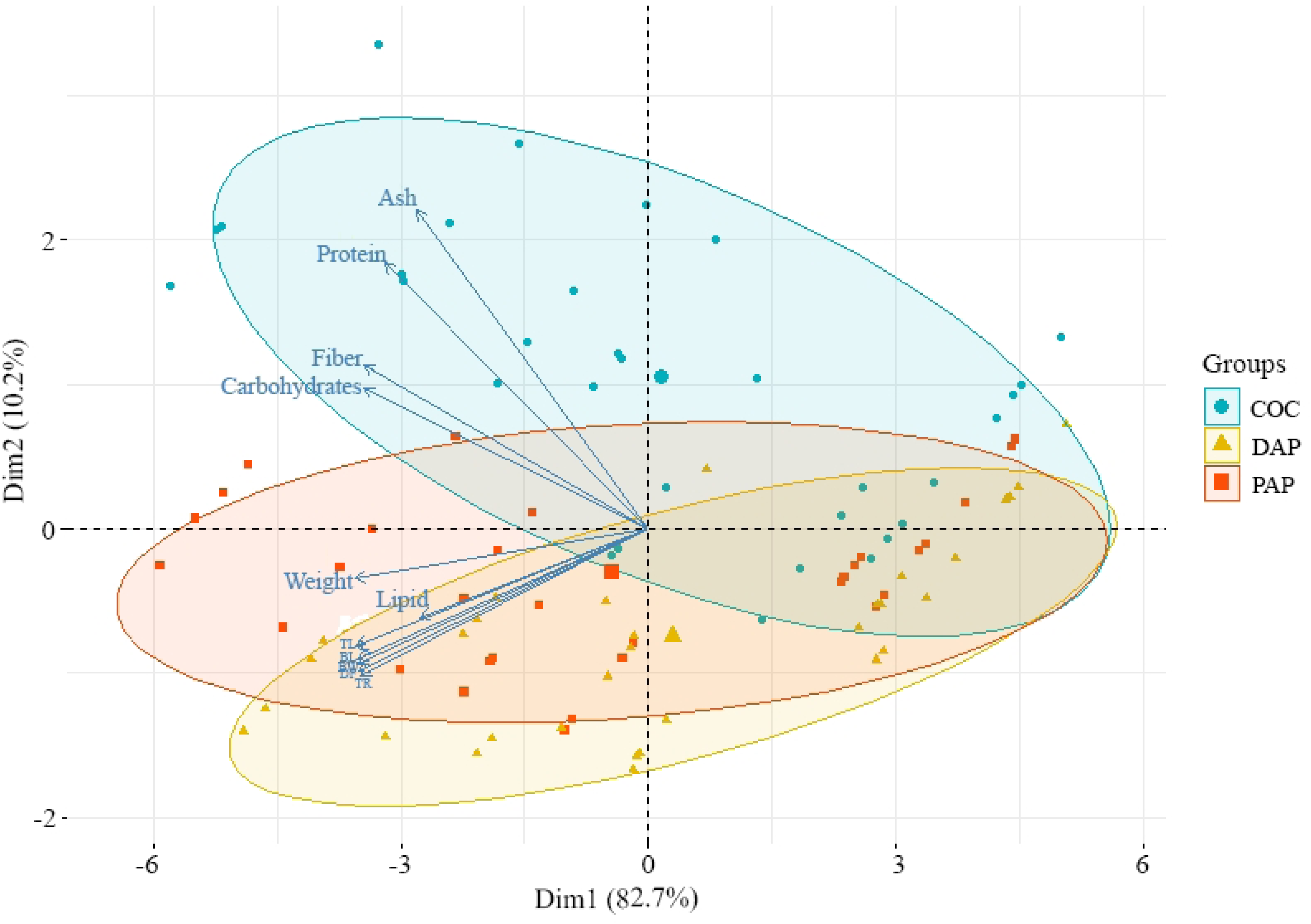
PCA correlating morphometric variables of juvenile Discus fish (*Symphysodon* sp.) and bromatological content of diets. COC: dry feed. DAP: live feed. PAP: wet feed.

## Discussion

The physicochemical parameters of the water in the aquariums remained within acceptable limits for this Amazonian cichlid species. Notably, nitrogenous waste products, such as ammonium (NH4) and nitrites (NO2), generated by uneaten feed and fish waste (Gilmour, 2022), remained at low concentrations. According to Kajimura et al. (2023), high concentrations of these compounds are toxic to fish, particularly Discus fish, which are highly sensitive to them (Ng et al., 2023).

Total dissolved solids (TDS) and conductivity represent the total mineral and salt content dissolved in the water (Cantillo & Corpus, 2018), classifying waters as either soft or hard (Diaz, 2018). In this study, the recorded values fell within ranges previously reported in the Amazon River: TDS (8.8–55 ppm) and conductivity (17.4–200 μS/cm) (Duncan & Fernández, 2010). Therefore, the water used in this study is classified as soft and corresponds to a tributary within the natural distribution range of *Symphysodon* sp. (Crampton, 2008).

The pH levels remained acidic throughout the experiment, optimal for Discus fish, which naturally thrive in habitats with a pH range of 6.0–6.8 (Duncan & Fernández, 2010) and in artificial environments with a pH between 6.8 and 7.0 (Pirhonen et al., 2012; Wen et al., 2018; Swain et al., 2020). Water temperature also remained within an optimal range. According to Pirhonen et al. (2012), Amazonian River waters typically range from 28°C to 29°C, enabling fish to consume and utilize feed more effectively, promoting adequate growth. Temperatures below 24°C or above 31°C negatively impact the development and survival of Discus fish (Bleher, 2006; Moncaleano & Riaño, 2018).

The selected feeds primarily consisted of plant-based ingredients, combined with some animal-based ingredients, aligning with the digestive physiology and dietary requirements of Discus fish (López-Rodríguez et al., 2019). As noted by Zabala-Camin (1996) and Crampton (2008), Discus fish have an omnivorous diet comprising algae and small aquatic invertebrates. This appropriate feed selection avoids the emergence of diseases and ensures optimal growth (Millan & Perez, 2017).

Juvenile Discus fish fed live feed combined with commercial feed (DAP) or wet feed combined with commercial feed (PAP) exhibited greater growth in length and weight than those fed commercial feed alone (COC). According to Koca et al. (2009) and Jiménez-Rojas et al. (2012), combining live feeds (e.g., *Enchytraeus buchholzi*, *Daphnia magna*) with commercial feed during the first 60 days in Angelfish (*Pterophyllum scalare*) led to greater weight gain. Live feeds are more uniformly distributed in the water column and stimulate predatory instincts (Velasco-Garzón & Gutiérrez-Espinosa, 2019). Similar results have been reported with other live organisms, such as Artemia and Tubifex, and nutritionally enriched water fleas, demonstrating optimal growth in cichlids (Jayalekshmi et al., 2017; Hudaidah et al., 2023; Mao et al., 2023).

For wet feeds, optimal growth in Discus fish (*Symphysodon aequifasciatus*) has been reported when combining prepared feeds with commercial feeds over a 60-day period (Santos et al., 2022). Wet feeds, due to their diverse ingredients, stimulate the secretion of digestive enzymes (Gonzales-Felix, 2010; Fuentes-Quesada et al., 2018). In this study, some of the ingredients used in the wet feed originated from the Amazon, including açaí (*Euterpe oleracea*), camu camu (*Myrciaria dubia*), and milpes (*Oenocarpus bataua*). This wet feed provided nutritional compounds found in their natural habitat. Furthermore, these Amazonian fruits contain vitamins, antioxidants, and antimicrobial properties (Portinho et al., 2012; Arellano et al., 2016; Yunis-Aguinaga et al., 2016), which contribute to cellular regeneration and fish growth.

The characteristics of live and wet feeds enable fish to feed and grow uniformly, minimizing hierarchical behavior by larger individuals that could negatively affect others (Luna-Figueroa et al., 2010). This behavior is often observed in Discus fish (Chong et al., 2000). Our results confirm that combining live and wet feeds with dry diets enhances feed conversion efficiency compared to single-feed types like dry feed. According to Luna-Figueroa et al. (2018) and Velasco-Garzón & Gutiérrez-Espinosa (2019), exclusive reliance on dry feeds may delay juvenile growth due to the lack of chemical and biological stimuli necessary for optimal growth.

The nutritional components of the diets are critical for fish growth and health (Velasco-Garzón & Gutiérrez-Espinosa, 2019). Our findings demonstrate that morphometric responses to the evaluated DAP and PAP diets are attributable to both the stimulatory effects of each feed type and their nutritional content. Proteins and fats were positively associated with juvenile growth. Notably, while proteins play a significant role in growth (Ribeiro et al., 2007; Gutiérrez-Espinosa et al., 2019), increasing their proportion does not necessarily accelerate growth. Instead, it may result in nutrient waste as fish fail to utilize excess protein efficiently (Mohanta & Subramanian, 2011; Velasco-Santamaria & Corredor-Santamaria, 2011). Other nutrients, such as lipids and carbohydrates, have a direct influence on growth by providing energy for essential metabolic processes (Prieto et al., 2008; Fracalossi & Cyrino, 2013). According to Sales & Janssens 2003, the presence of these components allows proteins to be used for tissue formation and repair rather than as an energy source, enhancing growth rates (Velasco-Garzón & Gutiérrez-Espinosa, 2019).

Lastly, the protein levels supplied in the diets aligned with the species’ dietary habits. Additionally, lipid- and carbohydrate-rich supplements, such as water fleas (*Moina macrocopa*) and wet feeds prepared with appropriate ingredients, are ideal for omnivorous species like Discus fish (*Symphysodon aequifasciatus*) (Burres et al., 2015). However, it is important to maintain low percentages of other nutrients, such as fiber (2–3%) and ash (8– 10%), to prevent digestive issues that could impair growth and health.

## Acknowledgments

The authors are thankful to the Servicio Nacional de Aprendizaje – Sena, Regional Caquetá and Centro de Investigación de la Biodiversidad Andino Amazónica – INBIANAM of the Universidad de la Amazonia for provide supplies, equipment and bromatological analyses to carry out this research.

